# A high-resolution map of non-crossover events reveals impacts of genetic diversity on mammalian meiotic recombination

**DOI:** 10.1101/428987

**Authors:** Ran Li, Emmanuelle Bitoun, Nicolas Altemose, Robert W. Davies, Benjamin Davies, Simon R. Myers

## Abstract

During meiotic recombination in most mammals, hundreds of programmed DNA Double-Strand Breaks (DSBs) occur across all chromosomes in each cell at sites bound by the protein PRDM9. Faithful DSB repair using the homologous chromosome is essential for fertility, yielding either non-crossovers, which are frequent but difficult to detect, or crossovers. In certain hybrid mice, high sequence divergence causes PRDM9 to bind each homologue at different sites, “asymmetrically”, and these mice exhibit meiotic failure and infertility, by unknown mechanisms. To investigate the impact of local sequence divergence on recombination, we intercrossed two mouse subspecies over five generations and deep-sequenced 119 offspring, whose high heterozygosity allowed detection of thousands of crossover and non-crossover events with unprecedented power and spatial resolution. Both crossovers and non-crossovers are strongly depleted at individual asymmetric sites, revealing that PRDM9 not only positions DSBs but also promotes their homologous repair by binding to the unbroken homologue at each site. Unexpectedly, we found that non-crossovers containing multiple mismatches repair by a different mechanism than single-mismatch sites, which undergo GC-biased gene conversion. These results demonstrate that local genetic diversity profoundly alters meiotic repair pathway decisions via at least two distinct mechanisms, impacting genome evolution and *Prdm9*-related hybrid infertility.

---

During meiosis, genetic information is exchanged between homologous chromosomes via the process of recombination. In mammals and other species, recombination is essential for the proper pairing of homologous chromosomes (synapsis) and their segregation into gametes, and together with mutation generates all genetic variation^1,2^. In many species, most recombination events cluster into small 1-2 kb regions of the genome, called recombination hotspots. In mice and humans, these hotspots are positioned mainly by PRDM9^3–8^, a zinc-finger protein that binds specific sequence motifs and deposits at least two histone modifications, H3K4me3 and H3K36me3^9,10^, on the surrounding nucleosomes. Double-Strand Breaks (DSBs) subsequently form near a small subset of PRDM9 binding sites in each cell^11^, and DSB processing results in single-stranded DNA decorated with the strand exchange proteins RAD51 and DMC1^7^. Each DSB can ultimately repair by homologous recombination in several ways (Fig. 1a). Because meiotic DSBs occur following replication of DNA, some DSBs can repair invisibly, using the sister chromatid as a repair template, although this process is disfavoured^12^. One exception is on the X chromosome in males, which has no homologue and instead repairs from its sister chromatid later in meiotic prophase^12^. The genetic features used for precise identification of the appropriate homologous DNA, to facilitate this repair, remain unknown. A minority of DSBs form crossovers (COs), involving reciprocal exchanges between homologues, while many more DSBs become non-crossovers (NCOs), in which a section of genetic material is copied (converted) from the homologue, without the donating chromosome being altered^13^.

**Fig. 1.**
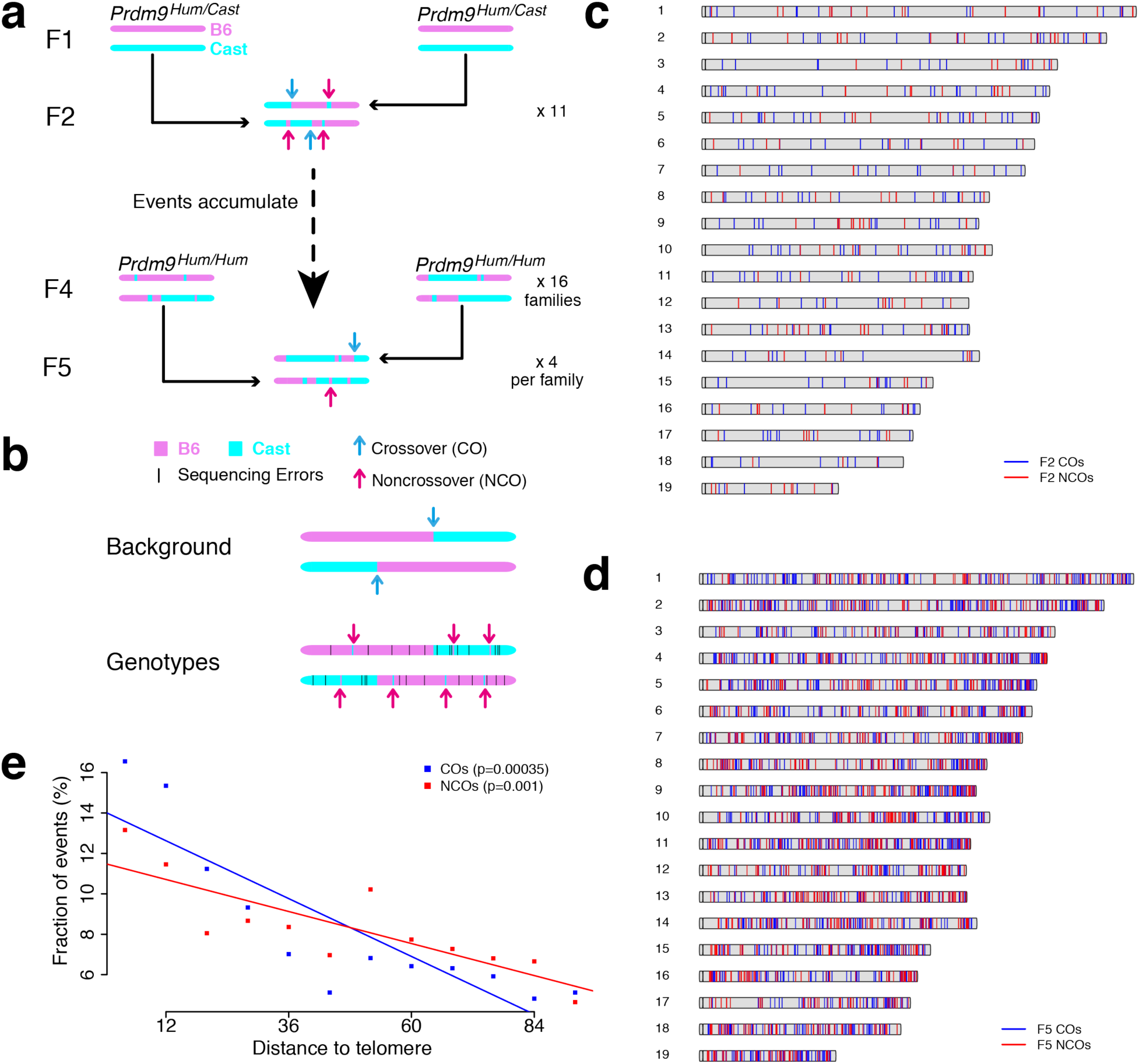
Study design and properties of CO and NCO events. **a** Study design. Arrows indicate locations of *de novo* CO and NCO events. **b** Detection of NCOs by comparing observed genotypes and background. **c**, **d** Distribution of identified COs and NCOs across autosomes from F2 (**c**) and F5 (**d**) animals. **e** Binning events by their distance to the telomere (x-axis). Full resolution available at: https://figshare.com/s/bf883f746fd676f1edb4

Although the role of recombination in shaping genetic variation is well understood, our understanding of possible effects in the reverse direction – of local genetic variation on NCO and CO event outcomes – remains very incomplete in mammals. Previous work^14–16^ has revealed impacts of genetic variation on the *formation* of DSBs, by altering PRDM9 binding properties. We also previously published evidence that genetic variation, by affecting PRDM9 binding, influences synapsis^17^, a process intimately connected to DSB repair. We found that the degree to which PRDM9 binding is “symmetric” – that is, whether PRDM9 binds both homologues equally well at each site – can influence synapsis and hybrid fertility^17^, with more asymmetric binding predicting reduced fertility in hybrid male mice. Moreover, individual asymmetric hotspots show elevated DMC1 ChIP-seq signals relative to H3K4me3, most consistent with the possibility that these DSBs take longer to repair^17^. One recent study^16^ reported that hotspots with high polymorphism rates, particularly those with asymmetry, show a stronger DMC1 signal compared to observed numbers of overlapping crossovers. This is consistent either with DSB repair delay, or with repression of COs within these hotspots. Despite this important progress, it therefore remains unclear how, or even if, local genetic variation might impact eventual recombination outcomes following DSB formation, or how this might differ for COs and NCOs and between sexes.

Furthermore, many fundamental questions remain about the process of non-crossover recombination itself, due to the difficulty of detecting NCO events. Previous studies in humans^18,19^ have revealed that in males, most (∼70%) NCO events occur within PRDM9-positioned recombination hotspots, and are short (<1 kb) and simple: they comprise contiguous tracts of converted SNPs, with no non-converted SNPs amongst them. In contrast, in females, a large number of “complex” NCO events, often extending over 1 kb, are seen. Long, complex human events are not strongly enriched in hotspots and show an association with maternal age^19^. Finally, human NCO events show a strong overall bias towards G/C bases (68%)^18,19^, as opposed to A/T bases^20–22^, occurring via an unknown mechanism. This phenomenon is thought to have driven regional differences in the GC-content of many species genome-wide^23–25^. Possible causes of this GC-bias include either subtle event initiation biases^24,26^, or heteroduplex DNA repair pathways^27^. However, simple models of heteroduplex DNA repair favouring G/C bases at mismatching bases are difficult to reconcile with the fact that most NCO events convert a contiguous set of SNPs, with no evidence of repair template switching. Moreover, not all SNPs in individually studied hotspots show GC bias^28^. It has been unclear to what extent these findings for humans might generalise to other species. Importantly, lack of power has prevented resolution thus far of fundamental questions about meiotic recombination, including any precise estimate of the length of underlying NCO tracts^18,19,29,30^, the total number of homologous recombination events per meiosis, and where NCO events position relative to PRDM9 binding sites and DSBs, although one recent study suggested a fairly broad distribution^27^.

To investigate links between genetic variation and repair outcomes, we mapped both CO and NCO events in mice, including mice humanized at *Prdm9*^17^, in both sexes. Critically, we also gathered complementary H3K4me3 and DMC1 ChIP-seq data (DMC1 data generated elsewhere^31^) in the male parental, or closely related, animals^14,16,17,19,31,32^ allowing us to analyse PRDM9 binding and DSB formation. Because H3K4me3 marks PRDM9 binding sites, at individual hotspots H3K4me3 signal strength approximates PRDM9 binding levels. DMC1 marks DSB sites prior to repair processing, so DMC1 signal strength at individual hotspots increases both with the rate at which DSBs occur, and the average time until these DSBs are repaired. Together, these data provide an unprecedented opportunity to investigate each step of meiotic recombination genome-wide and with high resolution, from PRDM9 binding, to DSB formation, to CO and NCO repair.

## Results

We identified both CO and NCO events in hybrids of two mouse strains: C57BL/6J, humanized at *Prdm9* (hereafter B6*^Hum^*, and predominantly of *Mus musculus domesticus* origin) and CAST/EiJ (hereafter CAST, predominantly of *Mus musculus domesticus* origin). Their high sequence divergence (0.7%; Methods) improves power to detect NCO events in offspring. B6*^Hum^* is identical to C57BL/6J except that the portion of the B6 *Prdm9* exon 10 encoding the DNA-binding zinc finger array has been replaced with the orthologous sequence from the human *PRDM9* B allele^17^, to produce a new allele we label *Prdm9^Hum^*, distinct from the *Prdm9^Cast^* allele possessed by CAST. The different *Prdm9* alleles allow us to distinguish the properties of *Prdm9^Cast^* and *Prdm9^Hum^* controlled recombination hotspots, with the humanized allele being of interest because it has not co-evolved with either mouse subspecies’ genome. We sequenced 11 F2 offspring of (B6xCAST)F1 mice (Fig. 1a), and after breeding for five generations in total to accumulate recombination events controlled by *Prdm9^Hum^*, we sequenced 72 (B6xCAST)F5-*Prdm9^Hum/Hum^* mice and their 36 F4 parents (Methods). We also gathered ChIP-seq data for both DMC1^31^ and H3K4me3 in testes from a male (B6xCAST)F1-*Prdm9^Hum/Cast^* mouse, allowing us to compare these to NCO/CO event outcomes. 23,748 DMC1 peaks correspond to DSB hotspots, and 63,050 PRDM9-dependent H3K4me3 peaks mark PRDM9 binding sites. For most peaks, we are able to determine which *Prdm9* allele controls them (Supplementary Note). Our data allow us to compare signatures of PRDM9 binding, DSB formation, and NCO/CO events, separately in both sexes.

To find both CO and NCO events, we developed and applied an HMM-based algorithm (Supplementary Note) to infer “background” states (B6/B6, B6/CAST and CAST/CAST) across the genome in each mouse to test potential gene conversions against (Fig. 1b). CO events correspond to background changes. SNPs with genotypes not matching their local background represent possible NCO events, but sequencing errors also mimic NCO events. Following careful filtering to exclude such errors (Methods and Supplementary Table 1) we identified 183 NCOs and 295 CO events on autosomes from the 11 F2 animals (Fig. 1c) and 1,392 NCOs and 2,205 COs in the F5 mice (Fig. 1d). This represents ∼3-fold more NCO events identified by direct sequencing, which avoids ascertainment biases, than in the largest previous mammalian study^19^, which was performed in humans - allowing inter-species comparisons. Sequencing-based validation of F2 events (Methods) estimated that 91% of the identified NCO events are real, while using simulations we estimate our power to identify those NCO events containing at least 1 SNP as 63% or above (Methods and Supplementary Fig. 1a, b). In the F5 mice, we were able to identify both *de novo* and parentally inherited NCO and CO events; and we were able to assign a subset to the maternal or paternal meiosis (Methods). NCO events can be assigned to a genetic background by determining whether they result from a DSB on the B6 or CAST chromosome.

### Overall event properties

NCO and CO events, as well as DMC1 and H3K4me3, show enrichment nearer to telomeres (Fig. 1e and Supplementary Fig. 1c). This is broadly similar to patterns observed in other mice^33–35^ and humans^15^, although COs show stronger enrichment than NCOs. Surprisingly and in contrast to events in human females, at least 99.4% of observed NCO events were “simple” and comprised contiguous tracts of converted SNPs, with no non-converted SNPs amongst them. Similarly, 99.4% of COs were simple background switches. This implies that complex NCOs are extremely unusual in mice. Moreover, we observed a very high overlap of both CO and NCO events with recombination hotspots, stronger than observed in humans^15,18,19^. In F2 mice (whose parents have the same F1 genetic background as the ChIP-seq samples), 96% of CO events and 92% of NCO events overlap ChIP-seq peaks (adjusted for false-positive NCO events and chance overlap; 84% unadjusted). This reduces only modestly in F5 mice, where only *Prdm9^Hum^*-controlled recombination hotspots are active (Supplementary Table 2), so hotspots identified in the heterozygous F1 mouse are still informative for meioses occurring in F4 mice. Thus, recombination hotspots identifiable by ChIP-seq account for essentially all recombination in mice, with little recombination in the remainder of the genome. Our findings confirm that female recombination mainly occurs in hotspots also active in male mice, in which our ChIP-seq data were gathered^36^.

NCO and CO events both occur in individual hotspots with probability approximately proportional to their estimated heat using either DMC1 or H3K4me3 signal strength (Fig. 2b, c): over 50% of all hotspot-associated F2 NCO or CO events occur in only the 4,000 hottest hotspots. Strong dominance of *Prdm9^Cast-^*controlled over *Prdm9^Hum^*-controlled hotspots is observed for both event types, and in our ChIP-seq data (Fig. 2d, e and Supplementary Fig. 2a, b). Because binding sites for the humanized allele have not experienced evolutionary hotspot erosion^14^, this phenomenon cannot explain the dominance of the *Prdm9^Cast^* allele. Instead, it could be due to a greater number of strong PRDM9^Cast^ binding targets genome-wide or possibly due to higher expression of *Prdm9^Cast^*.

**Fig. 2.**
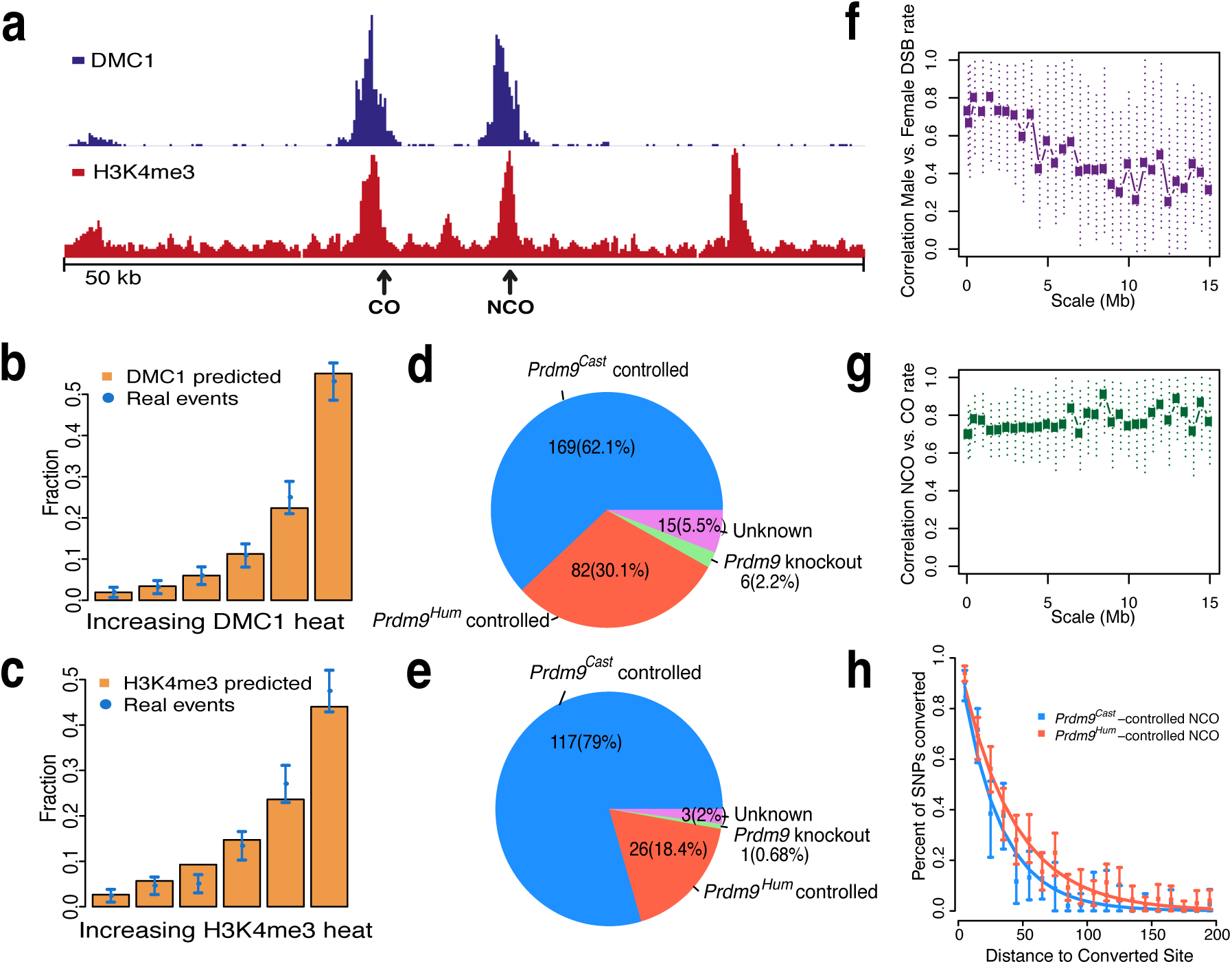
DMC1, H3K4me3, and *Prdm9* allele predict CO and NCO properties, and *Prdm9^Cast^* dominates *Prdm9^Hum^*. **a** DMC1 and H3K4me3 peaks in a 50 kb region on Chromosome 10, with single NCO and CO events overlapping these peaks. **b**, **c** DMC1 (**b**) and H3K4me3 (**c**) predict well where events occur. **d, e** Fraction of COs (**d**) and NCOs (**e**) controlled by the *Prdm9^Cast^* or *Prdm9^Hum^* alleles, overlapping hotspots in the *Prdm9* knockout mouse, or non-identifiable (Unknown). **f** Correlation of underlying recombination rates between females and males, for rates binned at different scales (x-axis); dotted lines show 95% CIs for true correlations. **g** As **f**, but showing correlations between (sex-averaged) NCO and CO rates at different scales. **h** Decay in probability that nearby SNPs are co-converted, with inter-SNP distance, conditional on a SNP being converted. Full resolution available at: https://figshare.com/s/bf883f746fd676f1edb4

After accounting for sampling variation (Methods), we estimated correlation between recombination rates at different scales (Fig. 2f, g and Supplementary Fig. 2c, d). This revealed sex differences, and strong (>70%) correlation between NCO and CO rates, although we also find very strong evidence that these events do differ in their positioning along the chromosome, especially at broad scales, and the NCO rate is much higher than the CO rate at all scales.

### Length, number, and positioning of NCO tracts

We leveraged the high SNP density in our system and large number of events to estimate the underlying NCO event tract lengths (accounting for the fact that if a NCO event does not contain a SNP, it is not observed; Methods), separately for hotspots controlled by *Prdm9^Cast^* and *Prdm9^Hum^*. The data show relatively good fits to an exponential distribution (Fig. 2h), but with significant differences in estimated mean NCO tract length (p=0.0018): 30 bp for *Prdm9^Cast^* (95% confidence interval (CI) 25-35 bp), and 41 bp for *Prdm9^Hum^* (35-48 bp CI). This is unexpected and implies that *Prdm9* alleles can differ in basic properties of how recombination events resolve. These tract length estimates are at the lowest end of the broad existing estimates for humans and mice^18,19,29,30^.

Previous studies using microscopy have reported 200-400 visible DMC1 foci marking individual DSB sites per meiosis in mice^29,37,38^. However, some of these DSBs, e.g. those occurring on the X-chromosome in males, might invisibly repair using the sister chromatid, which is present following replication. We directly estimated the total number of DSBs that repair from the homologous chromosome per meiosis. Using our tract length estimates (Methods), we inferred an average total of 300.5 DSBs (95% CI 258.5-370.5) per meiosis repairing using the homologue, 90% of these being NCOs^29,39,40^. Estimates are very similar in both F1 and F4 parents. Comparison with prior microscopy findings suggests that the majority of DSBs might be repaired via homologous chromosomes, rather than the sister chromatid^27,41^.

Both NCO and CO event centres distribute symmetrically around PRDM9 binding motifs that we identified within hotspots (Methods, Fig. 3a-d and Supplementary Fig. 3a-d). NCO events cluster very near to motifs (potentially overlapping them in 70% of cases; Fig. 3a, d), slightly less strongly than clustering of mapped DSBs^42^, but with a far tighter range than the DMC1 and H3K4me3 ChIP-seq signals, which identify single-stranded resection tracts around DSBs and histone methylation resulting from PRDM9 binding, respectively (Fig. 3e, f). CO events spread more broadly (Fig. 3b, c), consistent with previous studies^11,39,42^. Thus, NCO gene conversion appears restricted to sites very close to initiating DSBs themselves, and more distantly positioned NCOs^27^ appear to only occur rarely.

**Fig. 3.**
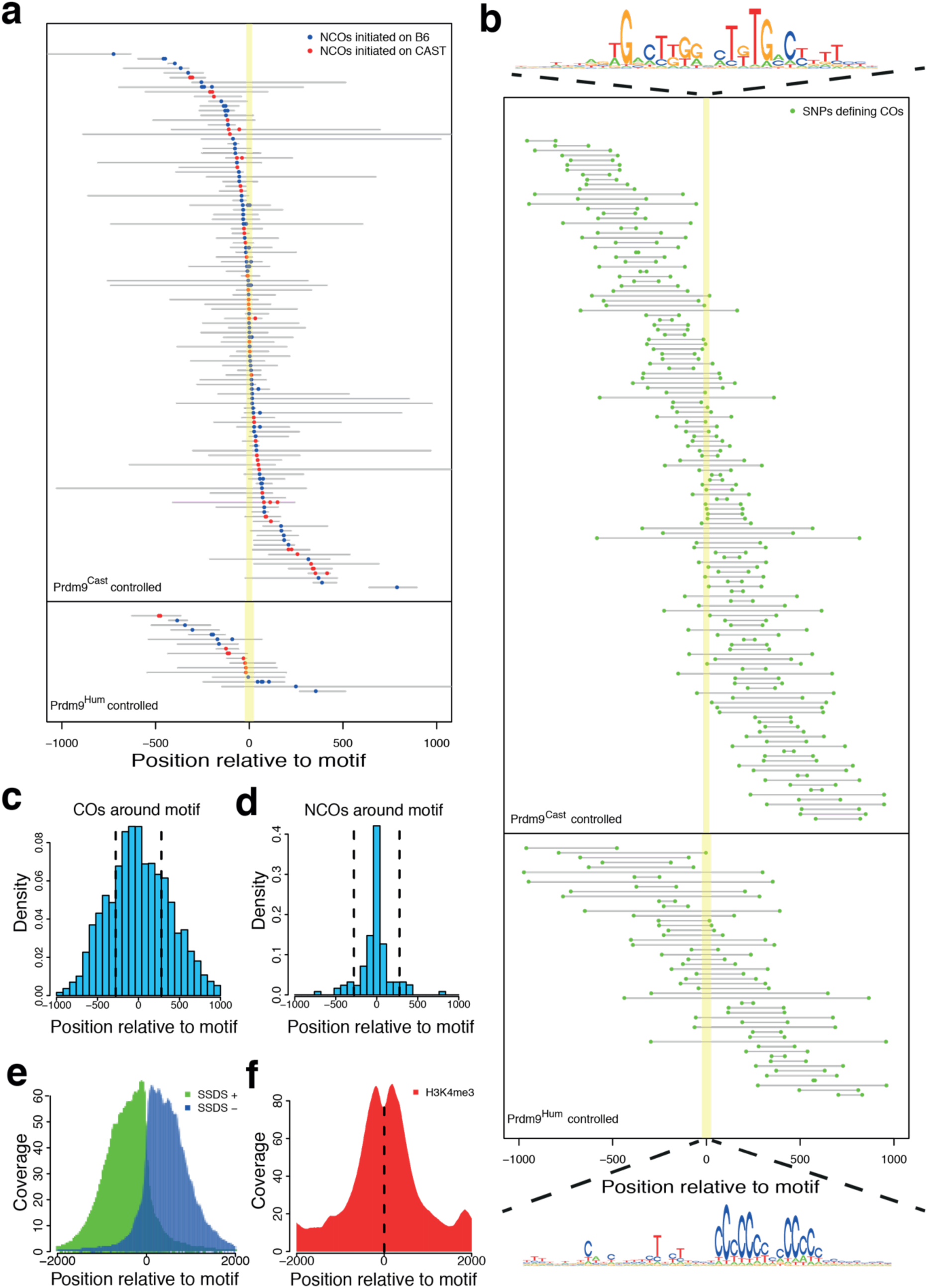
Position of NCOs, COs, and DMC1 and H3K4me3 peaks relative to PRDM9 binding motifs. **a** NCOs occurring within hotspots possessing robustly identified PRDM9 binding motifs. Coloured dots are converted SNPs and grey lines represent upper bound of converted tracts. Yellow shading indicates the identified PRDM9 binding target. **b** COs around PRDM9 binding motifs. Green dots are SNPs defining CO boundaries within grey delineating regions. COs that have large intervals (>2 kb) between the two defining SNPs are not shown in this plot. **c** Density of COs occurring around motifs. Bar height at each position is proportional to the probability that break point happens at this position and density in each bin is averaged across the positions. **d** Density of NCOs occurring around motifs. The distance between a NCO and motif is defined as the mid-point of minimal converted tract to the centre of the nearest identified hotspot motif. Distribution was normalised by SNP density in each bin to correct for increased power to see a NCO event where SNP density is high. **e, f** Mean DMC1 and H3K4me3 ChIP-seq read coverage around motifs, for the hotspots shown in **a** and **b**. For DMC1, we separated plus strand (SSDS+) and minus strand (SSDS-) reads. Note x-axis scale differs from **c** and **d**. Full resolution available at: https://figshare.com/s/bf883f746fd676f1edb4

### GC-biased gene conversion is controlled by SNP density and explains complex NCO and CO events

Our NCO events show strong evidence of AT-to-GC bias, though initially weaker than seen in humans^18^, for both *Prdm9^Cast^*-controlled (64%) and *Prdm9^Hum^*-controlled (60%) hotspots (p<2×10^−9^; Supplementary Table 3). We next focussed on NCO events within Prdm9^Hum-^controlled hotspots for further investigation, because the genomic GC-content has not evolved alongside this allele. We tested for a difference in NCO tracts containing a single SNP with those containing multiple SNPs (Fig. 4a). Surprisingly, this revealed GC-bias to occur exclusively in single-SNP NCO tracts, which show a near-identical GC-bias (68%) in both males and females (Supplementary Table 4). In complete contrast, no bias (p=0.92) is seen for all multiple-SNP tracts combined, and the difference relative to single-SNP tracts is highly significant (p=1.1×10^−7^). We exactly replicated this finding (p=5.6×10^−4^) in NCO events within hotspots in humans^19^, in both males and females (Supplementary Table 4), so it represents a conserved phenomenon across these mammals. GC-bias strength is unaltered even if DSBs happen only on one homologue (Supplementary Fig. 4a and Supplementary Table 4), implying a mechanism driven by heteroduplex repair^27^ rather than DSB formation^24,26^.

**Fig. 4.**
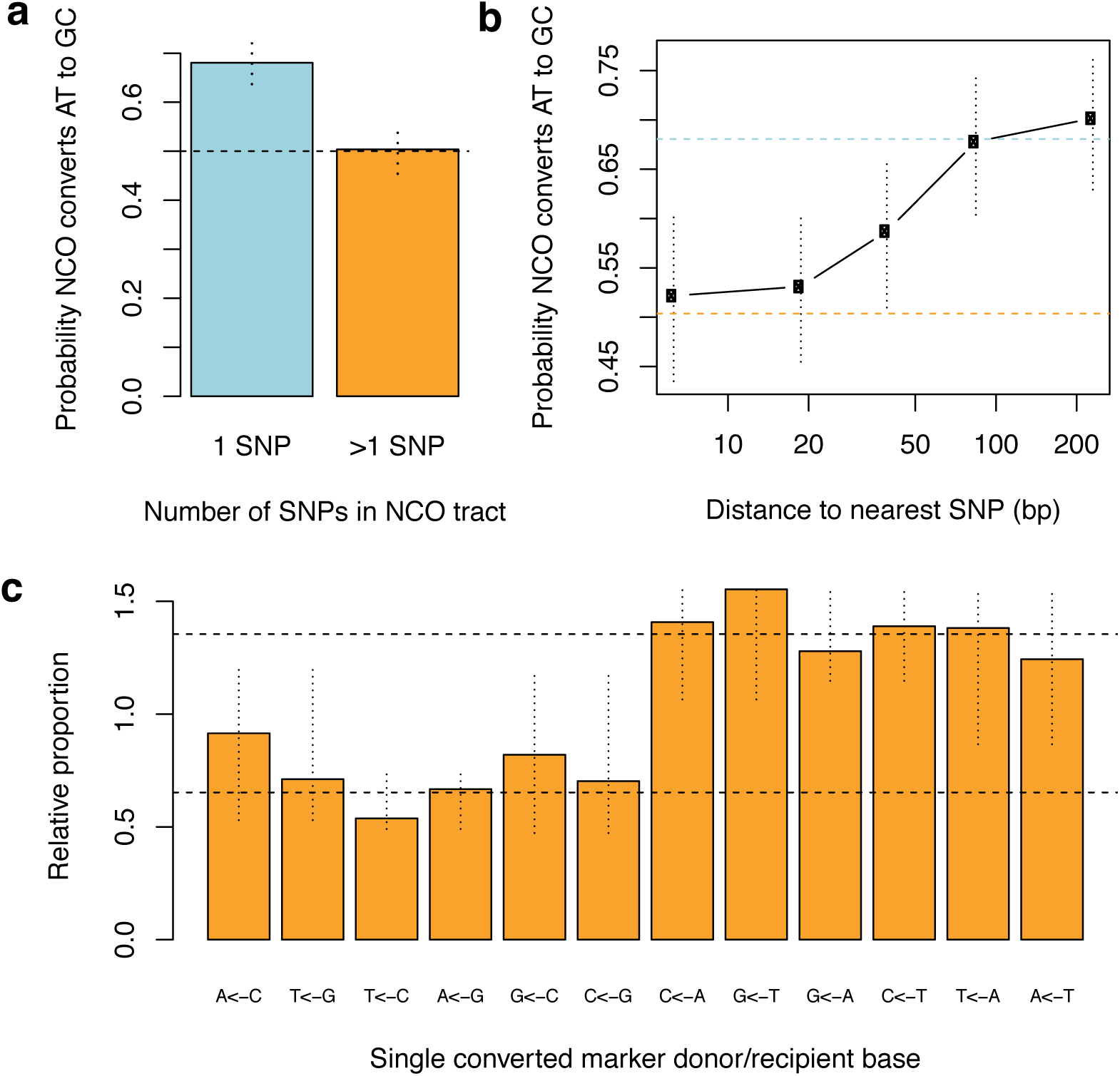
GC-biased gene conversion is absent in multi-SNP NCO tracts, and SNPs nearby other SNPs show no detectable GC-bias. **a** AT-to-GC conversion in single-SNP vs multiple-SNP tracts. **b** GC-bias in groups of converted SNPs, binned according to their distance to the nearest SNP. **c** For each of the 12 possible combinations of NCO donor/recipient alleles (x-axis; e.g. A<-C converts recipient C to donor A), the proportion of observed single-SNP NCOs of that type is plotted, relative to the corresponding proportion for the nearest non-converted markers, which lack GC-bias. Vertical lines: 95% CIs after pooling strand-equivalent pairs. Horizontal dotted lines: mean relative proportions for NCO events whose recipient types are G/C or A/T respectively. Full resolution available at: https://figshare.com/s/bf883f746fd676f1edb4

A restriction of GC-bias to single-SNP tracts might reflect either some GC-biased process preventing longer events occurring, or a direct impact of the number of SNPs within heteroduplex DNA on whether GC-bias occurs. To distinguish between these possibilities, we stratified SNPs by distance to their nearest SNP and measured their GC-bias if they fell within NCO events (Fig. 4b). Strikingly, SNPs near to other SNPs, and therefore almost always co-converted with them, show no GC-bias evidence. Conversely SNPs further than typical NCO tract lengths, >100 bp from the nearest SNP, show the ∼68% bias observed in humans, in whom SNP density is much lower^18,19^. This implies that local genetic diversity itself influences GC-biased gene conversion at NCOs, and therefore there must be at least two distinct processes operating to repair heteroduplex stretches formed at DSBs, one which is strongly GC-biased, and another which dominates when multiple mismatches exist, and shows no GC-bias.

To further characterise GC-bias, we estimated conversion rates of different types of SNP in the donor and recipient chromosomes at single-SNP NCO sites (Fig. 4c and Supplementary Note). We normalised these relative to their conversion rates in multi-SNP events (Supplementary Fig. 3b), or to flanking SNP composition (Fig. 4c), both of which show no GC-bias and gave near-identical results. The simplest model which can explain the data is if there are two distinct conversion rates, with observed NCO rates lower if the *recipient* chromosome (i.e. the homologue in which the DSB occurs) carries a G or a C, and higher if the recipient carries an A or a T. For example, G/C transversions appear to convert at the lower rate. This could be explained by a model where a GC-biased process can resolve heteroduplex DNA in favour of the recipient chromosome, if it carries a G and/or C base – effectively “blocking” conversion of that base. If so, higher local heterozygosity, which disrupts this process, would be expected to actually *increase* local NCO rates.

Interestingly, we do not observe a consistent GC-bias for CO events, which are accompanied by long conversion tracts of ∼500 bp in size^30^. However, we did observe a very small number of “complex” events, incorporating non-converted markers surrounded by converted markers, and resulting from the same meiosis. We hypothesised that these might result from occasional operation of the GC-biased process. If so, the above results suggest that complex events might result from “blocking” of conversion of particular markers where the recipient chromosome carries a G or C base. This motivates examining the non-converted markers surrounded by converted markers within complex CO and NCO events. We observed a total of 12 such markers within NCO events and 7 within CO events. Remarkably, for 18 of these 19 cases the recipient chromosome carries a G or C base (p= 7.6×10^−5^ by 2-sided binomial test).

Therefore, in our mice essentially all of the complex NCO and CO events we observe can be explained in terms of the action of a GC-biased process which normally only operates within single-SNP conversion tracts. A recent study of one human hotspot^43^ found a similar GC-bias of 87-100% for complex male CO events, so it seems likely this process operates across species. Moreover, the bias of nearly 100% towards the recipient carrying a G/C, compared to the ∼68% bias of all single-SNP NCO events, might suggest that non-biased heteroduplex repair occurs even in some tracts containing only a single heteroduplex site. For example, the bias might only impact either G or C recipient bases, but not both, which would cap the bias at NCO sites to (at most) 67%, very close to the observed fraction.

### Hotspots where the homologue is not bound by PRDM9 show reduced homologous recombination

For NCO events, we can identify on which homologue the underlying DSB occurred. In the F2 mice, we observed a bias: 60% of the observed NCOs were initiated on the B6 background (p<10^−3^). This is due to the behaviour of the *Prdm9^Cast^* allele, which accounts for 80% of observed NCO events (Fig. 2e), and which shows a strong preference for binding to the B6 background (Supplementary Fig. 5a; 66% of NCOs, p<10^−3^), explained by evolutionary hotspot erosion of CAST-controlled hotspots on the CAST genetic background^14,16^. As expected, the *Prdm9^Hum^* allele binds and initiates recombination events equally on both backgrounds (Supplementary Fig. 5a, p=0.63). The fraction of NCOs initiating on the B6 background correlates highly with the fraction of DMC1 and H3K4me3 ChIP-seq signals originating from that background (Supplementary Fig. 5b, e). Because the ChIP-seq data only reflect male meiosis but observed NCO events originate from both males and females, this high correlation implies similar hotspot behaviour in both sexes. The similar increases in PRDM9 binding, DSB formation, and NCO formation on the B6 chromosome imply that no strong compensation mechanism acts to equalise the number of DSBs or recombination events on different homologues, although weaker compensation that we lack power to detect might occur.

Recombination hotspots can be separated into “asymmetric” cases where DSBs occur mainly on one homologous chromosome, and “symmetric” cases where DSBs occur equally on both homologues. Using H3K4me3 and DMC1 ChIP-seq data, we estimated the fraction of PRDM9 binding and DSB formation on the B6 vs. CAST chromosome in each hotspot^17^. Among the most asymmetric hotspots, we observed SNPs or indel polymorphisms within 96% of identified motifs overall (Supplementary Fig. 5f), implying their asymmetry is almost always driven by sequence changes disrupting PRDM9 binding on one homologue. Therefore, asymmetric binding, and hotspots, are conserved between the sexes and between F2 and F5 animals.

As in other hybrid mice^17^, we observed that the ratio of DMC1 to H3K4me3 ChIP-seq signal increases roughly two-fold at asymmetric hotspots compared to symmetric hotspots (Supplementary Fig. 5g), a fact most easily explained by delayed repair of DSBs forming at positions whose homologue is not bound by PRDM9^17^. We tested whether such asymmetry might also disrupt DSB repair processing, by quantifying asymmetry (Methods) and measuring the numbers of NCO and CO events actually occurring in asymmetric vs. symmetric hotspots, relative to their expectations according to DMC1 and H3K4me3 signal strength (Methods and Supplementary Note).

H3K4me3 signal strength approximates PRDM9 binding levels, while DMC1 levels at individual hotspots reflect numbers of DSBs (as well as repair timing). As we found across events overall (Fig. 2), with all else being equal we expect any two groups of hotspots that are matched to have the same total DMC1 or H3K4me3 signal to also have similar numbers of CO and NCO events. However, when we grouped *Prdm9^Hum^*-controlled hotspots according to their (a)symmetry we instead observed a strong depletion of both NCO and CO events in the most asymmetric hotspots (p=10^−27^ and p=10^−23^, respectively, after controlling for factors influencing power; Fig. 5), whether DMC1 or H3K4me3 was used. We replicated this signal for *Prdm9^Cast^*; for both males and females; and for *de novo* and inherited events in F5 mice, as well as events in F2 mice (Supplementary Note and Supplementary Fig. 6), so this is a general property of asymmetric hotspots in both sexes. Because the *Prdm9^Hum^* allele in particular did not co-evolve alongside the mouse genome, asymmetric hotspots controlled by this allele reflect chance genetic variation disrupting PRDM9 binding sites on one homologue or the other, implying a mechanistic impact of asymmetry on recombination independent of hotspot erosion or other evolutionary forces.

**Fig. 5.**
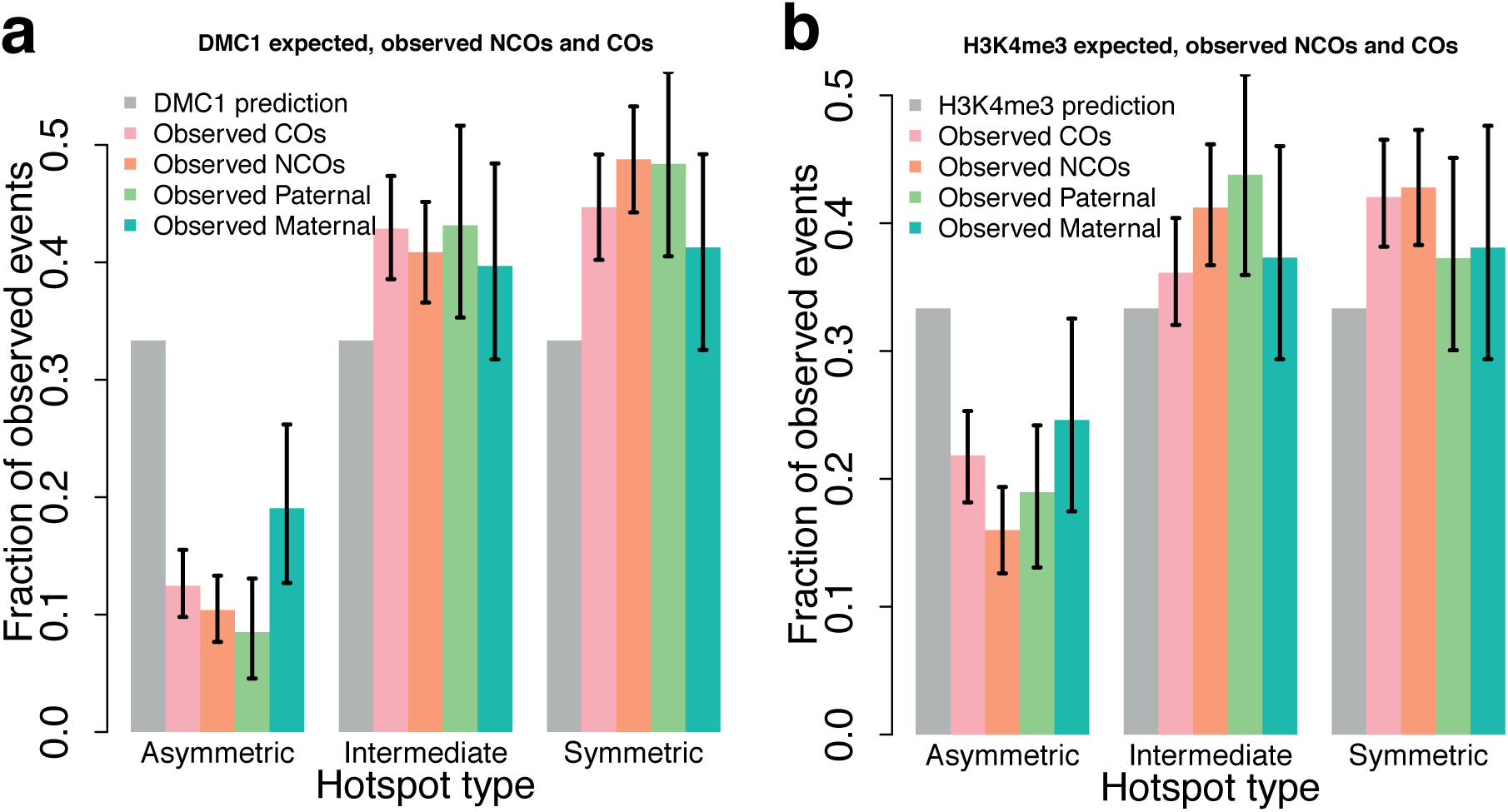
COs and NCOs are depleted in asymmetric hotspots in both males and females. **a** Human-controlled DMC1 hotspots were separated into 3 bins (asymmetric, intermediate, symmetric) according to symmetry, so that each bin contains the same number of predicted events according to DMC1 heat. Grey bars show the DMC1-predicted expected fraction of events in each bin. The four coloured bars (vertical lines: 95% CIs) show the observed fraction of (sampled) F5 *de novo* events: COs, NCOs, and paternal or maternal recombination events. **b** As **a**, except predicted events were defined using H3K4me3. Full resolution available at: https://figshare.com/s/bf883f746fd676f1edb4

Importantly, we found that this homologous recombination deficiency is driven by PRDM9 binding asymmetry alone, rather than SNP diversity elsewhere within hotspots (Methods). Furthermore, for DSBs occurring on the less-bound chromosome of asymmetric hotspots, we found that NCO events occur at the expected rate for symmetric hotspots (Supplementary Note). This implies that when DSBs occur at asymmetric hotspots on the *more* frequently bound chromosome, the resulting lack of observed NCO or CO recombination events must be specifically driven by a lack of PRDM9 binding to its homologue, rather than any impact of diversity *per se*, even within PRDM9 binding motifs.

Because H3K4me3 reflects the level of PRDM9 binding, the lack of homologous recombination at asymmetric hotspots might in principle be explained if DSBs occur less often at these sites. However, this seems impossible to reconcile with the dramatic *excess* DMC1 signal observed at asymmetric hotspots for a given level of H3K4me3 signal. Instead, our results can be explained if, at DSB sites whose homologue is not bound by PRDM9, repair sometimes takes place using the sister chromatid rather than the homologous chromosome. This must occur in both sexes. Sister-based repair can explain the lack of observable COs and NCOs in such cases, and the identical magnitude of depletion of each type of event, implying that asymmetric hotspots fundamentally disrupt interactions between homologous chromosomes. This implicates PRDM9 binding motifs as one genetic “signal” allowing identification of DNA on the homologue, to facilitate homologous repair.

Interestingly, sister chromatid repair is thought to operate on meiotic DSBs on the X chromosome in males^12^, which, similar to autosomal DSBs in asymmetric hotspots, exhibit a very strong increase in DMC1 signal relative to H3K4me3 signal, probably owing to the late timing of their repair^17^. If repair via the sister chromatid occurs later in general, this would explain the observed elevation of DMC1 signal relative to H3K4me3 at asymmetric hotspots.

## Discussion

Although complex NCO events are common in human females^18,19^, with an incidence increasing with maternal age, we found that such events are near absent in mice. We suggest that this difference may reflect differences in the timespan of dictyate arrest, which occurs before the completion of female recombination, and lasts decades in humans vs. months in mice. These findings support the hypothesis that complex NCO events in humans might reflect the repair of non-programmed DNA damage occurring over time, consistent with the fact that they are different in other ways: they mainly occur outside PRDM9 hotspots^19^, are often >1 kb long, and show GC-bias regardless of their size.

Our results indicate a sex-averaged NCO rate in mice carrying humanized *Prdm9* of around 10^−6^ per base. This is strikingly below human estimates of around 4.1×10^−6^ and 7.7×10^−6^ in males and females respectively^19^, meaning humans show even greater increases in the NCO:CO ratio relative to mice – for unknown reasons. Nonetheless, our minimum estimates of total DSB counts in mice are consistent with previous microscopy studies, suggesting that most DSBs in mice repair using the homologous chromosome, at least those not found in asymmetric hotspots or impacted by GC-biased gene conversion. Speculatively, perhaps a higher rate of NCO events in humans may aid synapsis by providing more potential inter-homologue interaction sites during meiosis. However, an elevation in homologous recombination events also increases the potential for mispairing at some sites, with consequences including diseases caused by non-allelic homologous recombination.

Using DMC1^31^ and H3K4me3 ChIP-seq data, one can infer that DSBs form at a small subset^11^ of PRDM9 binding sites in each cell on average proportionally to the rate of PRDM9 binding to each site^17^. However, we find that the processing, repair and eventual recombination outcomes at each PRDM9 binding site all depend strongly on the sequence of the homologue. In this study, we revealed for the first time that both CO and NCO events are depleted, in both sexes, at “asymmetric” recombination hotspots where one homologue is not strongly bound by PRDM9 (primarily due to mutations in the PRDM9 binding motif on the homologue; Fig. 5). We also showed that this reduction in homologous recombination cannot be explained by an increase in genetic diversity alone, as has been suggested^16^; only nearby SNPs that abolish PRDM9 binding symmetry have any effect on the CO or NCO rate. Furthermore, for DSBs occurring on the less-bound homologue in asymmetric hotspots, we found that NCOs occur at the expected rate for symmetric hotspots. This proves that homologous recombination outcomes at each DSB are strongly influenced by PRDM9 binding to the homologue and/or modifying its histones. That is, when the homologous site is not bound by PRDM9, the DSB is less able to repair by homologous recombination, which is critical for synapsis and fertility.

Others have recently demonstrated a depletion of COs at asymmetric recombination hotspots relative to DMC1 signal strength^16^. However, their data could not discriminate between the following causal possibilities: (i) genetic diversity *per se* was responsible; (ii) DMC1 elevation through repair delay entirely drove the signal; or (iii) these sites were preferentially repaired instead by NCO recombination. Here, we have ruled out these possibilities by demonstrating an equal depletion of CO and NCO recombination events exclusively at asymmetrically bound sites, even relative to expected event numbers from PRDM9 binding data (H3K4me3). This in turn allows us to confidently infer that some process downstream of DSB formation, but upstream of the CO vs. NCO repair decision, is disrupted at asymmetric sites. We suggest that this disrupted process may be homology search, rather than mismatch repair^16^ (which would impact NCO events occurring on both homologues similarly). One hypothesis is that PRDM9 binding and/or chromatin marks on the unbroken homologue assist the homology search machinery with the challenging task of finding the correct homologous template for repair; without this assistance, homology search and homologous recombination are impaired at asymmetric sites. Together with previous work^16^, our data indicate that suppression of recombination is a general property of asymmetric hotspots. This mechanism can explain the wider asynapsis and infertility seen in male mice where asymmetric hotspots predominate^17,44^, although additional factors must act to explain sex differences in hybrid fertility.

We also confirmed our previous finding that when DSBs form at asymmetric hotspots, they show a two-fold excess of DMC1 signal relative to H3K4me3 signal on the more strongly bound homologue, which is consistent with delayed DSB repair^17^. Taking into account their elevated DMC1 signals, asymmetric hotspots behave oddly, by showing a combination of (at least in males) slower DSB repair, and an inability to engage with their homologues for this repair, via either CO or NCO, in both sexes. These phenomena could be explained by a model in which many DSBs in asymmetric hotspots fail to interact with their homologue and are instead repaired late from the sister chromatid. That is, asymmetric hotspots may behave like DSBs on the X chromosome in males, which repair late and from the sister chromatid, and show excess DMC1 signal^12,17^. Other models would involve either more or fewer DSBs occurring at asymmetric hotspots and seem unlikely, because they would require strong pairing of homologues prior to DSB formation in order to distinguish symmetric and asymmetric binding sites.

In addition to altering PRDM9 binding symmetry and the resulting recombination outcomes, local genetic differences between homologous chromosomes can also alter the process of GC-biased gene conversion (gcBGC) at NCO sites. We confirmed gcBGC operates downstream of DSB formation, and this implies it must act on repair of heteroduplex DNA with mismatching bases, formed during DSB repair towards recombination. We found that in both humans and mice, within hotspots gcBGC acts almost exclusively on potential conversion tracts containing only a single SNP (i.e. mismatch), with essentially identical bias in each species (68% of NCOs convert A/T to G/C)^18,19^. This single-SNP preference can explain why most multi-SNP NCO tracts are simple stretches of markers without “cherry-picking” of markers converted towards GC. Because heteroduplex DNA is expected to form for all possible NCO and CO tracts containing SNPs, not just those containing single SNPs, our results imply more than one pathway for heteroduplex repair (Fig. 6). In the first, gcBGC acts to favour the strand on which the DSB occurs: if this strand carries a G or C at the SNP, conversion is prevented from occurring. We calculated that such a process would need to block gene conversion at G/C recipient sites 53% of the time to account for the observed 68% overall GC-bias in observed events (Fig. 6 and Supplementary Note). Almost all observed complex NCOs and COs, though very rare, appear to be explained by this GC-bias preventing conversion of individual markers, with the background on which the DSB occurred carrying a G or C base in 95% of such non-converted markers we observed. Similar behaviour was observed in a study of male COs within a single human hotspot^45^. Among suggested drivers of mammalian gcBGC^20^, these properties of strong (almost 100%) base-specific and strand-specific biases, and operating at very fine scales (single SNPs), appear most consistent with the action of base excision repair (BER) rather than mismatch repair (MMR) proteins.

**Fig. 6.**
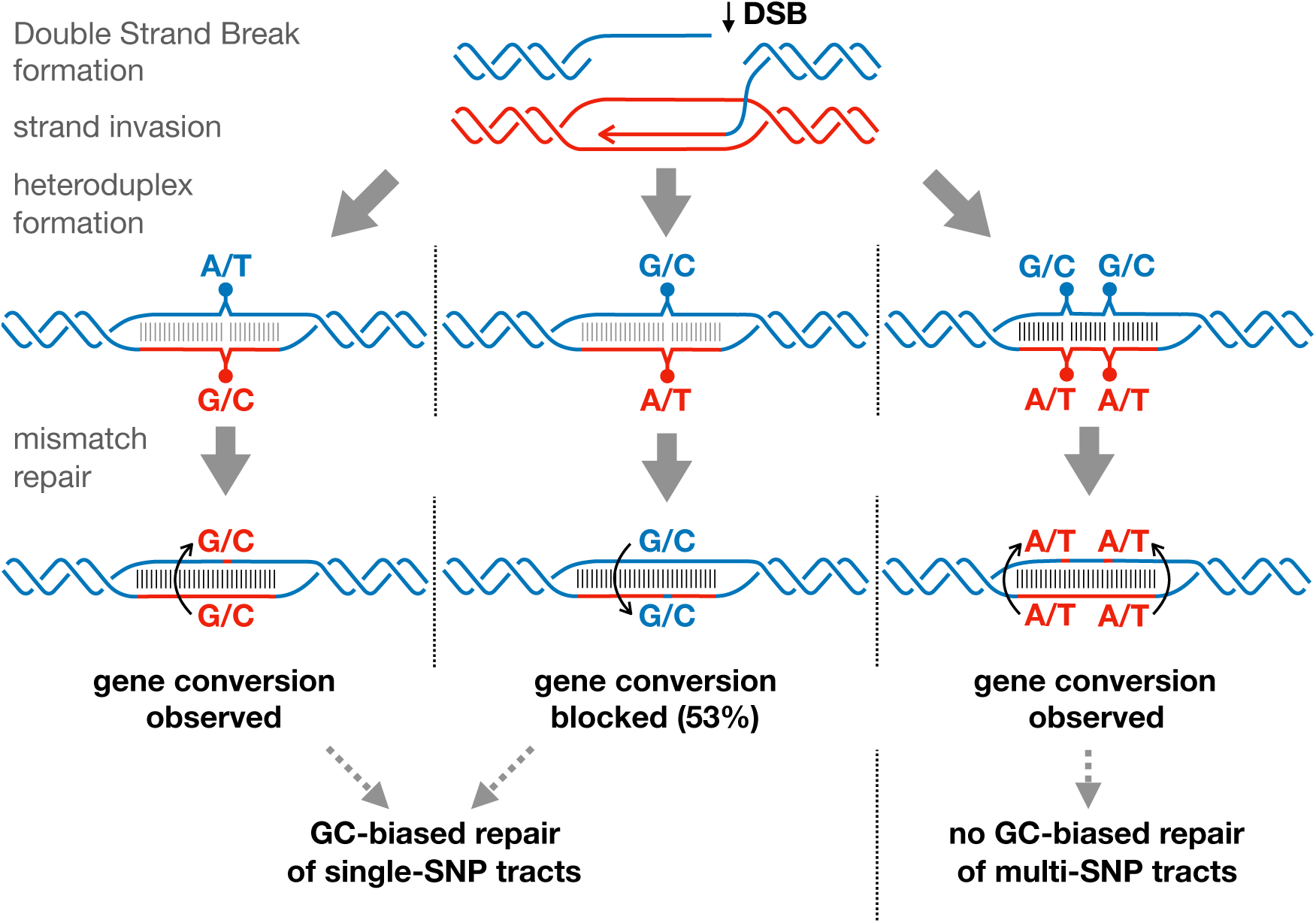
Model explaining influence of local genetic diversity on non-crossover mismatch repair pathway choice. Three possible gene conversion tracts are depicted, differing in the number and type of heteroduplex mismatch sites on the recipient chromosome (blue). In the first case (left), a single A/T site on the recipient chromosome is converted in a strand-biased manner (perhaps by MMR) to the allele of the donor chromosome, regardless of donor base type (red). When the recipient chromosome contains a G/C at a single mismatch site (middle), a different repair mechanism (perhaps BER) operates 53% of the time and blocks gene conversion (Supplementary Note). The sum of these two effects can explain why 68% of observed gene conversions are converted to G/C. When a second mismatch is present nearby (right), repair reverts to a strand-biased mechanism, and no GC-bias is observed, except in rare complex NCO events. Full resolution available at: https://figshare.com/s/bf883f746fd676f1edb4

The alternative, non-GC-biased repair pathway of heteroduplex DNA instead can act on multiple-SNP stretches, and must show a strand bias – this time favouring the *incoming* strand, copied from the homologous chromosome on which the DSB did *not* occur (Fig. 6). Otherwise, if heteroduplex mismatch repair had no strand bias, half of potential NCOs would repair invisibly, and so to account for our estimate of the NCO rate (∼274 NCOs per meiosis) there would need to be twice as many DSBs per meiosis (∼600), which is far outside the range of previous experimental observations^29,39,40^. Moreover, although resolution of heteroduplex DNA towards the broken chromosome within potential NCO events would be invisible, at 19 observed CO events within highly (>95%) asymmetric hotspots containing a mutation within their PRDM9 motif, we observe transmission of the “cold” binding site allele to offspring in 95% of cases. Therefore, for those longer conversion tracts within CO events at least, heteroduplex repair appears overwhelmingly biased towards the unbroken homologue. If this mechanism for resolving heteroduplex mismatches is also used for single-SNP stretches in some cases, this would explain why gcBGC appears weaker for NCO events in general, compared to complex NCO and CO events. We suggest that this non-GC-biased process, impacting longer stretches including multiple SNPs, is consistent with properties of MMR proteins, several of which are known to be essential for meiosis in mice^45,46^. If these hypotheses are correct, it is interesting that BER and MMR appear able to favour different strands.

Together, these gcBCG results imply that SNP density within hotspots influences recombination events downstream of DSB formation. For example, the same SNP will show different conversion rates and biases in different individuals, depending on nearby heterozygosity patterns in those individuals. Interestingly, this predicts a slightly higher NCO gene conversion rate in more diverse than less diverse regions. Another unexpected influence on NCO events, again impacting downstream of DSB formation, is the PRDM9 allele, with PRDM9^Hum^-controlled NCOs having an average length 11 bp (37%) longer than PRDM9^Cast^-controlled NCOs. It is unclear whether this reflects PRDM9 binding directly, or some indirect impact, e.g. how PRDM9 binds relative to nucleosome positions.

The effects of genetic diversity on PRDM9 binding and DSB formation are increasingly well understood^14–17^. However, there has been little prior knowledge of how genetic diversity can influence recombination after DSB formation (though the converse—how recombination affects genetic diversity—is highly studied). Here we have uncovered the first examples of local genetic diversity affecting both the ability to undergo homologous repair (due to polymorphisms altering PRDM9 binding symmetry), and the decision to repair via GC-biased gene conversion (depending on the number and spacing of local sequence polymorphisms), both downstream of DSB formation. Interestingly, our results predict that the reduction of homologous recombination at asymmetric hotspots should somewhat mitigate hotspot erosion caused by the over-transmission of alleles that disrupt PRDM9 binding^14,17^. Further work will need to investigate the exact mechanisms by which PRDM9 binding to the unbroken homologue promotes homologous repair, or the exact repair pathways that can explain the observed GC-biased or strand-biased NCO recombination outcomes.

## Methods

### Mouse breeding, library preparation and sequencing

CAST/Eij (CAST) mice were sourced from MRC Harwell (UK). The C57BL/6J (B6) line humanized at the *Prdm9* zinc-finger array (B6^Hum^) was generated previously^17^. Breeding of CAST and B6^Hum^ mice (F0) was carried out in both directions (using females and males of each type) to generate (B6xCAST)F1 hybrid, heterozygous offspring. To study properties of the humanized *Prdm9* allele, we genotyped as previously^17^ and selected 26 F2 mice of each sex homozygous for humanized *Prdm9*. We crossed these animals and their offspring for 3 further generations and selected 72 F5 offspring, comprising 2 of each sex from each of 18 pairs of F4 parents. One B6^Hum^ mouse, one CAST mouse, 11 F2 mice and all 18 F4/F5 families (36 F4 parents and 72 F5 offspring) were subjected to whole genome sequencing. Genomic DNA was extracted from spleen using the DNAeasy Blood and Tissue Kit (Qiagen), according to the manufacturer’s instructions. Libraries were prepared by the Oxford Genomics Centre at the Wellcome Centre for Human Genetics (Oxford, UK) using established Illumina protocols (with a Nextera DNA Library Prep Kit).

We sequenced to obtain coverage of ∼10x for the F0/F4 mice, and 20x for the F2 and F5 mice via the Illumina Hiseq2500 (F0 and 4 F2 mice) or Hiseq4000 platforms (remaining mice). Genomic DNA was fragmented to an average size of 500 bp and incorporated into libraries using established Illumina paired-end protocols (Nextera DNA Library Prep). Sequencing reads were aligned to mm10 using BWA^48^ (v. 0.7.0) followed by Stampy^49^ (v. 1.0.23, option bamkeepgoodreads). We used Picard tools (v. 1.115) (http://broadinstitute.github.io/picard) to merge bam files from different lanes for the same sample and identify duplicate reads. GenomeAnalysisTK-3.3-0 (GATK) was used for local Indel realignment followed by base quality score recalibration, and variant calling, using known Indel targets and SNPs between B6 and CAST from the Mouse Genome Project (MGPv4) data^50^. We filtered variants using GATK’s Variant Quality Score Recalibrator (VQSR), employing the set of variants present on the Affymetrix Mouse Diversity Genotyping Array as a set of true positive variation^51^. We used the annotations “HRun”, “HaplotypeScore”, “DP”, “QD”, “FS”, “MQ”, “MQRankSum”, and “ReadPosRankSum” to train VQSR, and a sensitivity threshold of 90% for the true positive set to define the set of newly genotyped sites that passed VQSR filtration. To remove potential hidden heterozygous sites within the F0 individuals, we removed all variants not genotyped as matching the homozygous reference allele in B6, or the homozygous alternative allele for CAST from MGPv4^50^. We obtained 13,946,562 and 13,940,079 reliable autosomal SNPs from the F2 and F5 samples, or roughly one SNP for about every 170 bp, which were used for downstream analysis.

### Identifying unique NCO and CO events

Using the HMM method described in the Supplementary Note to define a background state (homozygous CAST background, heterozygous, or homozygous B6 background) along the genome in each mouse, we identified state changes as CO events. Autosomal genotypes in F2 and F5 mice conflicting with their background were investigated as potential NCO events, but mainly represented sequencing errors. The region chr6:37000000-56000000 (mm10) was removed since it was observed to be not fully inbred in the F0 founders. We filtered to remove false positive sites (Supplementary Table 1), apparently heterozygous sites in the F0 mice, false heterozygous calls exhibiting unequal numbers of reads supporting the two alleles, false homozygous calls due to low read depth, and others. The number of potential converted sites dropped greatly e.g. from 863,082 SNPs potentially converted to 183 distinct identified NCOs within the F2 mice.

For COs and NCOs identified in F5 animals, they were treated as inherited if the parents carried an identical event, and otherwise *de novo*. We used a previously described HMM algorithm^52^ to identify parent-of-origin in *de novo* COs (those occurring in the germ cells of the F4 parents). For *de novo* NCOs, we were only able to confidently assign parental origin when one of the parents was heterozygous at the converted sites, while the other was homozygous. In this case, the NCO must be inherited from the heterozygous parent.

We removed duplicate inherited CO and NCO events, yielding a set of unique events for downstream analyses. Of 1,575 observed NCO events, only 9 were “complex” and involved background switching within the event. Of 1,116 observed *de novo* CO events from F2 and F5 animals, 7 were complex e.g. a CO accompanied by a NCO event.

### NCO validation by Sanger sequencing

To validate a subset of NCO events detected in F2 mice (19 within and 9 outside a hotspot), we PCR-amplified short regions (around 200 bp) overlapping the identified NCO sites using genomic DNA from the 2 F0 mice, the F2 mouse carrying the NCO, and up to 3 other related and/or unrelated F2 mice, using standard conditions (cycling conditions and primer sequences available upon request). PCR products were purified using the QIAquick PCR Purification Kit (Qiagen) and analysed by direct Sanger sequencing (Source Bioscience, UK). Sequence data comparison and analysis was carried out using Chromas LITE (version 2.1.1). By comparison of genotypes, we identified true NCOs vs. false positives. This confirmed all 19 NCOs overlapping a hotspot (100%), and of 9 non-hotspot cases, 4 were confirmed (44%). This suggests almost all hotspot-overlapping NCO events are likely real, and a higher false positive rate in NCOs away from hotspots. Given that 84.2% of the F2 NCO events overlap hotspots (Supplementary Table 2), we estimate an overall fraction of validated detected NCO events as 0.842 + 0.44 x 0.158, i.e. 91.1%.

### Estimating power to identify NCOs

To estimate the power of our method to detect NCO events of varying tract lengths, we simulated NCOs with different mean tract lengths and ran our pipeline for identifying NCO events, including our filters. Because F2 events are controlled by both *Prdm9^Hum^* and *Prdm9^Cast^* and F5 *de novo* events are controlled by *Prdm9^Hum^* alone, we performed two sets of simulations by using data from 11 F2 samples and 72 F5 samples. Because most recombination events overlap hotspots, we simulated NCOs in hotspot regions. For each mean tract length, we sampled 2,000 hotspots with probabilities proportional to their H3K4me3 enrichment. Within each hotspot, we sampled the centre of the NCO tract according to the distribution of NCOs around PRDM9 motifs after correcting for SNP density, and we sampled its tract length from an exponential distribution with a pre-defined mean tract length (which we varied from 10 to 100 bp with step size 10 and from 150 to 300 bp with step size 50). Sampled NCO tracts containing 0 SNPs were not counted as potentially detectable. Across these 2,000 tracts, different animals possessed different ancestral backgrounds. For each tract in each animal, we checked if any of the other animals had a different ancestral background consistent with a gene conversion event in the first animal. If so, we sampled such a “donor” mouse (other events were ignored). We copied the sequencing information corresponding to the converted sites from the donor mouse, such as the allele depth, and we copied the sequencing information for the background from the recipient, such as mate-pair information. Then, we applied the same filters to this simulated sequencing data at each sampled tract. We calculated our power by dividing the total number of simulated tracts left after filtering by the total number of simulated tracts overlapping at least one SNP (Supplementary Figure 1a, b).

### H3K4me3 ChIP-seq

We performed ChIP-seq against H3K4me3 in testes from an 8-week-old male (B6xCAST)F1-*Prdm9^Hum/Cast^* mouse C57BL/6J-*Prdm9^Hum/Hum^* mother, CAST/Eij father) as previously described^17^ with several important modifications that increased ChIP stringency (noted here). Lysis was performed in 1% SDS lysis buffer (1% SDS, 10 mM EDTA, 50 mM Tris pH 8.0, 2x protease inhibitors). Sonication was performed in a Bioruptor Twin sonication bath (Diagenode) at 4°C for three 5-minute periods of 30s on, 30s off at high power. Sonicated lysates were diluted 1:10 in IP wash buffer (containing 500 mM LiCl) instead of dilution buffer for antibody incubation. This yielded roughly 1 ng of ChIP DNA per testis. ChIP and total chromatin DNA samples were sequenced in multiplexed paired-end Illumina HiSeq2500 libraries (rapid run), yielding 63-71 million 51-bp read pairs per replicate after filtering (one ChIP replicate per testis plus one input sample). Sequencing reads were processed and peaks were called as described in our previous work^17,53^. Haplotype assignment of ChIP signal and removal of PRDM9-independent H3K4me3 peaks were performed as described^17^. The percentage of ChIP-seq read pairs originating from signal (as opposed to background) was estimated to be 87.4%, a significant improvement over our prior, less stringent, experimental method (which yielded 62-71% of read pairs from signal)^17^.

### DMC1 ChIP-seq

DMC1 ChIP-seq data were generated elsewhere^31^. Briefly, single-stranded DNA sequencing (SSDS) DMC1 ChIP-seq was performed as described previously^54^, using testes from a male (B6xCAST)F1*^Prdm9^* ^*Hum/Cast*^ mouse. ChIP and total chromatin DNA samples were sequenced in multiplexed paired-end Illumina HiSeq2500 libraries (rapid run), yielding 252 million 51-bp read pairs. We processed the data following the algorithm provided by Khil *et al*. 2012 to map the reads to mm10 and obtain type I reads^54^. We then called DMC1 peaks as previously^17^. We defined NCO and CO events as occurring within hotspots if they were less than 1 kb away from either a DMC1 peak or an H3K4me3 peak (covering 4% of the genome).

### Testing for correlation of different recombination-related phenotypes at chosen scales

We assumed that given an underlying vector of (binned) mean values *W_k_* along the genome, the k^th^ recombination-related quantity (number of observed recombination or NCO events in various classes), *N_ik_*, follows a Poisson distribution with mean *W_ik_*, in interval *i*. The *W_k_* means vary along the genome and represent the underlying recombination rate parameters; this model is accurate provided (as is likely to be case) for a single meiosis, the number of expected events in each bin is small. Then the variance

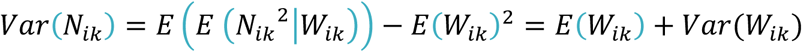

This enabled estimation of the variation in recombination rate along the genome, using the usual standard estimates of the mean and variance of the number of events, across bins genome-wide:

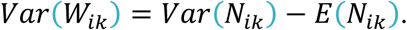

Further, the covariance

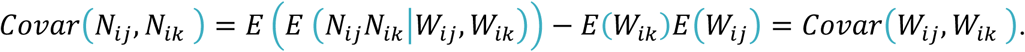

Combining these results enabled estimation of the underlying correlation between *W_j_* and *W_k_* along the genome based on properties only of the observed Poisson counts *N_j_* and *N_k_*:

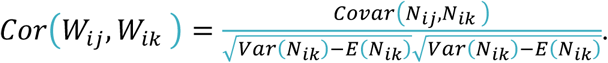

The quantities in the above equation are all estimated in the usual way using standard estimates of mean and variance from the observed vectors of counts. At any interval size scale, we bootstrap re-sampled (10,000 times) the resulting disjoint intervals of the genome, to compute CIs for the estimator.

### Motif Analysis

We used a Bayesian, *ab initio* motif finding algorithm to identify motifs within DSB hotspots^17,53^. For each DSB hotspot that is controlled by *Prdm9^Cast^*, a 1,000 bp sequence (centred on the hotspot centre) was extracted from the reference sequence (mm10). *Ab initio* motif identification was performed on the centre 600-bp sequences from the top 1,000 hotspots (ranked by DMC1 heat) that contained no bases overlapping annotated repeats. Motif calling proceeded in two stages: seeding motif identification, and motif refinement. Each seeding motif was obtained by first counting all 10-mers present in all input sequences, and from the top 50 most frequently occurring 10-mers, the one with the greatest over-representation in the central 300 bp of each peak sequence was chosen. This seeding 10-mer was then refined for 50 iterations as described in Davies *et al.*, 2016. This refined motif was then force-called on the full set of the hotspots (without filtering) by re-running the refinement algorithm, providing a probability of motif occurrence within each hotspot, and also identifying the most likely motif location in each case. This motif was reported for each peak, along with position and strand. We did the same for DSB hotspots controlled by *Prdm9^Hum^* and a 48-bp human motif was identified. We identified distinct sequence motifs, and their locations, within 97% of hotspots controlled by *Prdm9^Cast^* and 74% of hotspots controlled by *Prdm9^Hum^* (Supplementary Fig. 3a)^14,17,19,55^.

We used the SNPs generated as described above to determine whether each motif contains a SNP within its span. The distance from a motif to an event was defined as the distance from the centre of the motif to the nearest converted marker (lower bound for NCOs), or zero if a converted marker fell within the motif itself. We associated events <1 kb from a motif with that motif-containing hotspot.

### Estimation of NCO tract length for human-controlled and CAST-controlled events

To estimate NCO tract length, we assumed the converted tract follows an exponential distribution with rate parameter λ, where 1/λ is the mean tract length. While exponential tract lengths are not a fully accurate model, we can view this as a summary of tract properties, estimating the probability of co-conversion of pairs of markers as the distance between them increases. We computed a composite likelihood function for our NCOs and estimated λ via maximal likelihood. Specifically, for each converted site, viewing this site as a “focal” site, we examined the SNPs nearby and recorded for each SNP its distance from the focal SNP, and whether that SNP was also converted. If the SNP was also converted, then it was still in the gene conversion tract, otherwise it was not. Using this approach allowed our approach to be independent of SNP density, because we conditioned on SNP positions in our analysis. The probability that a SNP nearby a converted site is also converted is

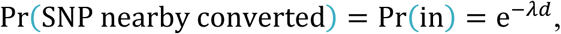

where *d* is the distance from the nearby SNP to the converted site. The probability that a SNP nearby a converted site is not in the tract is 1-Pr(in). All the NCOs are independent so we can multiple these probabilities for each SNP in the windows to get the (composite) likelihood of the data:

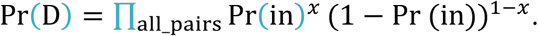

Here *x*=1 if the SNP nearby is also converted and *x*=0 otherwise. By maximising the likelihood using grid search for 1/λ from 1 to 1000 with step 0.1, we gained an estimate of tract length. Because pairs of SNPs are not in fact independent, this is not a true likelihood (though the resulting estimator is statistically consistent as the number of independent conversion events increases), and so to estimate uncertainty in the resulting estimates, we utilised bootstrapping of NCO events.

To perform bootstraps, we separated autosomal genomes into 258 non-overlapping 10 Mb blocks (the last block in each chromosome is shorter than 10 Mb). We re-sampled 258 blocks with replacement, where the probability of sampling each block is proportional to the length of that block, and from the resulting bootstrapped set of NCOs, re-estimated tract length via the same procedure. CIs were calculated from a total of 10,000 bootstraps. We implemented this procedure for two sets of NCO events; those overlapping human-controlled and those overlapping CAST-controlled hotspots, respectively.

### Hotspot symmetry estimates

Sequence differences between the CAST and B6 genomes allowed us to quantify the fraction of ChIP-seq signal (either DMC1 or H3K4me3), coming from the B6 and CAST chromosomes. This also allowed us to determine whether individual hotspots in these hybrids were ‘symmetric’, with DSBs occurring equally on both chromosomes, or ‘asymmetric’, with a preference towards either the CAST or B6 chromosome.

Using SNPs distinguishing the B6 and CAST genomes, each type I read pair from a hybrid DSB library (DMC1 ChIP-seq) was assigned to one of the categories ‘B6’, ‘CAST’, ‘unclassified’ or ‘uninformative’ as in^17^, replacing PWD with CAST. For each DSB hotspot, the B6 cutting ratio was then computed as the fraction of ‘B6’ reads mapped within 1 kb of the hotspot centre, over the sum of ‘B6’ and ‘CAST’ reads in that region. We followed a similar approach for H3K4me3 ChIP-seq, further correcting for background signal as in^17^. For both DMC1 and H3K4me3, we required ≥10 informative reads to define the B6 cutting ratio.

To order hotspots based on their symmetry, if the fraction of cuts estimated on the B6 and CAST chromosome were *x* and 1-*x*, respectively, we defined the overall hotspot “symmetry” as 4*x*(1-*x*), which ranges from 0 for hotspots with events completely on one chromosome to 1 for hotspots with events occurring equally on both chromosomes^17^. We obtained additional results for events initiating on a known homologue by using “homologous heat”, defined as *xh*, where *h* is the estimated total heat of the hotspot, for events initiating on the CAST chromosome, and (1-*x*)*h* for events initiating on the CAST chromosome (Supplementary Note). Separate estimates of hotspot symmetry and homologous heat may be obtained from both H3K4me3 and DMC1 ChIP-Seq data, for the same collection of hotspots. Because the H3K4me3 homologous heat captures how well the homologous chromosome is bound by PRDM9, it may be of stronger direct interest; however, homologous heat is only directly identifiable for NCO events, whose initiating homologue is known. For CO events, to be conservative (avoiding assumptions regarding conversion tracts to estimate homologous heat), we mainly used hotspot symmetry instead of homologous heat. For Supplementary Fig. 6 we used average homologous heat, defined as 2*hx*(1-*x*), which averages homologous heat over the strand an event occurs on.

### Estimation of the fraction of asymmetric/symmetric hotspots containing a motif-disrupting variant

To estimate the proportion of hotspots of different levels of initiation on B6/CAST chromosomes containing SNPs within their PRDM9 binding motifs, we filtered to include only hotspots containing a clear motif (posterior probability >0.99), and at least 20 informative reads in our DMC1 data in order to accurately estimate the proportion of reads from B6, and 5 sequencing reads from each homologue covering the motif region, to provide power to identify variants if present. Supplementary Fig. 5f shows the fraction of hotspots in each binned level of initiation on the B6 chromosome containing a SNP or Indel (using GATK prior to VQSR, or Platypus). 96% of identified highly asymmetric hotspots (B6 initiation <5% or >95% and P<10^−10^ for binomial test of asymmetry) contained such a polymorphism.

### Testing whether asymmetry or SNP density affects the resolution of recombination events

We fitted a generalised linear regression model to discern whether hotspot asymmetry or local SNP density better predicts variation of CO and NCO rates depending on genetic variation. For each hotspot containing an identified PRDM9 binding motif, we produced a binary response vector indicating whether an overlapping CO event occurred and fit a binomial generalised linear model. As model predictors, we used:

i. The symmetry of the hotspot
ii. The log-transformed ‘heat’ of the hotspot measured by H3K4me3 (the H3K4me3 heat is incremented by a small value 0.0001 as there are a few hotspots with zero estimated heat)
iii. SNP densities around the PRDM9 binding motif at different scales (±100 bp, ±500 bp, ±800 bp)

We then tested each coefficient for significance, conditional on the others. We separated the analysis for *Prdm9*^Cast^-controlled COs (all generated in the meiosis from F1 where there are two different *Prdm9* alleles) and *de novo Prdm9^Hum^*-controlled COs (all generated in the meiosis from F4 where there is only one type of *Prdm9* allele) to eliminate any effects of competition between the two alleles. Conditional on the heat of H3K4me3 and hotspot symmetry, SNP density has no significant effect on where COs happen (p-values from all three scales >0.08) while both heat and symmetry of hotspots have significant positive effects on CO events conditional on SNP density (p<0.05).

For NCO events, we performed a similar analysis, except that we corrected for power to detect NCOs by re-sampling the above hotspots according to the weight generated as described in the Supplementary Note section “Rejection sampling for COs and NCOs, construction of Fig. 5 and Supplementary Fig. 6, and testing for impacts of asymmetry on event resolution”. We note that some hotspots appeared several times after rejection sampling. Again, all scales showed no significant effect of SNP density conditional on the heat of H3K4me3 and hotspot symmetry (p>0.2). For *Prdm9^Hum^*-controlled NCOs, results show that the heat of H3K4me3 and symmetry of hotspots have significant positive effects on NCOs conditional on SNP density (p<0.003). Results from *Prdm9^Cast^*-controlled NCOs also suggest positive effects on prediction of NCOs, but p-values do not reach significance due to the smaller number of these events (<0.2). We discuss the weaker effect of symmetry for *Prdm9^Cast^*-controlled NCOs in the Supplementary Note.

### Data availability

The datasets generated and analysed during the current study will be made available in public repositories (GEO and SRA) prior to publication. The H3K4me3 ChIP-seq data are currently available with GEO accession GSE119727.

### Code availability

The computer code developed for the analysis of the datasets in the current study will be made available in Github prior to publication.

### Ethical compliance

All experiments involving research animals received local ethical review approval from the University of Oxford Animal Welfare and Ethical Review Body (Clinical Medicine board) and were carried out in accordance with the UK Home Office Animals (Scientific Procedures) Act 1986.

## Acknowledgments

We thank the High-Throughput Genomics Group at the Wellcome Centre for Human Genetics (funded by Wellcome Trust grant reference 203141/Z/16/Z) for the generation of sequencing data. We also thank Gang Zhang and Anjali Gupta Hinch for generating and providing the DMC1 ChIP-seq data for the (B6xCAST)F1-*Prdm9^Hum/Cast^* mouse. Funding: This work was supported by two Wellcome Trust Investigator Awards to S.R.M. (098387/Z/12/Z and 212284/Z/18/Z); N.A. is a Howard Hughes Medical Institute Gilliam Fellow.

## Author contributions

S.R.M. designed the study; B.D. generated the humanized B6 mouse; E.B., N.A., and B.D. bred the mice (F0-F2: N.A. and B.D.; F3-F5: E.B.); E.B. and N.A. prepared the genomic DNA samples (6F0/F2: N.A and 115 F2/F4/F5: E.B); E.B. performed NCO event validation; N.A. performed H3K4me3 ChIP-seq and data processing; R.L. analysed the data; R.W.D. contributed SNP data; R.L., E.B., N.A. and S.R.M prepared the manuscript; B.D. critically reviewed the manuscript.

## Materials & Correspondence

requests should be addressed to S.R.M.

